# DNA-SIP and repeated isolation corroborate *Variovorax* as a key organism in maintaining the genetic memory for linuron biodegradation in an agricultural soil

**DOI:** 10.1101/2020.11.15.383406

**Authors:** Harry Lerner, Başak Öztürk, Anja B. Dohrmann, Joice Thomas, Kathleen Marchal, René De Mot, Wim Dehaen, Christoph C. Tebbe, Dirk Springael

## Abstract

The frequent exposure of agricultural soils to pesticides often leads to microbial adaptation, including the development of dedicated microbial populations that utilize the pesticide compound as a carbon and energy source. Soil from an agricultural field in Halen (Belgium) with a history of linuron exposure has been studied for its linuron-degrading bacterial populations at two time points over the past decade and *Variovorax* was appointed as a key linuron degrader. Like most studies on pesticide degradation, these studies relied on isolates that were retrieved through bias-prone enrichment procedures and therefore might not represent the *in situ* active pesticide-degrading populations. In this study, we revisited the Halen field and applied, in addition to enrichment-based isolation, DNA stable isotope probing (DNA-SIP), to identify the *in situ* linuron degrading bacteria. DNA-SIP unambiguously linked *Variovorax* and its linuron catabolic genes to linuron dissipation, likely through synergistic cooperation between two species. Additionally, two linuron mineralizing *Variovorax* isolates were obtained with high 16S rRNA gene sequence similarity to strains isolated from the same field a decade earlier. The results confirm *Variovorax* as the *in situ* degrader of linuron in the studied agricultural field and corroborate the genus as key in the maintenance of a robust genetic memory regarding linuron degradation functionality in the examined field.

## Introduction

Pesticides ensure sufficient crop yields and their application is an essential component of the global food supply chain. However, these compounds are often persistent and can affect environmental and human health, thus demanding strict regulation of their authorisation and use (Bolognesi, 2003; Kristoffersen *et al.* 2008; Li and Jennings, 2017; Nicolopoulou-Stamati *et al.* 2016). Along with its chemical and physical properties, biodegradability is a pivotal attribute of a pesticide, determining its persistence and hence its fate and associated risks (Arias-Estévez *et al.* 2008; Fenner *et al.* 2013; Karpouzas *et al.* 2016). The biodegradability of pesticides relies largely on the diverse bacterial community that inhabits the soil environment and therefore understanding the ecology of pesticide-degrading microbes is crucial for fully comprehending the fate and environmental impact of pesticide compounds. In recent years it has been demonstrated that soils, when frequently exposed to a pesticide, can develop a strong biodegradation capacity and activity for that pesticide due to the selection and proliferation of dedicated pesticide-degrading bacterial populations that use the compound as a source of energy and carbon or for acquisition of other essential nutrients, often leading to its mineralization (Imfeld and Vuilleumier 2012; Pesce *et al.* 2013; Jiang *et al.* 2017). However, our knowledge of the *in situ* degraders of pesticides and their dynamics in soil environments, including agricultural soils, is still scarce and largely depends on knowledge derived from pesticide-degrading isolates obtained from enrichment cultures (ECs). For instance, independently isolated degraders of a particular pesticide often belong to the same bacterial genus, indicating a form of specialization among bacteria for acquiring pesticide degrading abilities (Lal *et al.* 2006; Simonsen *et al.* 2006; Breugelmans *et al.* 2008; Nguyen *et al.* 2014; Yan *et al.* 2016). However, whether those micro-organisms are actually responsible for *in situ* pesticide biodegradation is currently unknown, considering that enrichment is prone to biases, for instance due to preferential growth or horizontal gene transfer during the enrichment process. The well-studied microbiology and microbial ecology underlying the biodegradation of the phenylurea herbicide linuron (3-(3,4-dichlorophenyl)-1-methoxy-1-methyl urea) continues to be a relevant model system in this context (Hussain *et al.* 2015).

Linuron is a non-selective pre-emergence herbicide affecting the photosystem-II of broad-leafed weeds. The compound is banned in the European Union and other countries due to accumulating evidence of human health risks (European Comission; Government of India), but is still in use in others like the USA and Canada (Government of the USA; Government of Canada). Linuron exposed soils, including agricultural soils and soils near pesticide production plants, have been the source for enriching and isolating various linuron-degrading bacterial strains and consortia from several geographic locations such as Belgium, Denmark, and China (Dejonghe *et al.* 2003; Sørensen *et al.* 2005; Breugelmans *et al.* 2007; Zhang *et al.* 2018, 2020). These isolates allowed the elucidation of the linuron catabolic pathway, which starts with the hydrolysis of linuron to 3,4-dichloroaniline (DCA), and continues with the degradation of DCA to chlorocatechol, which is further degraded to oxoadipate and channeled into the Krebs cycle (Dejonghe *et al.* 2003). Members of the *Variovorax* genus appear to be especially relevant, since many linuron-degrading isolates belong to this genus. They act either as single strains performing all necessary steps to grow on the compound, or as part of consortia in which they perform at least the crucial step of linuron hydrolysis (Dejonghe *et al.* 2003; Sørensen *et al.* 2005; Breugelmans *et al.* 2007; Satsuma 2010). These linuron-degrading *Variovorax* isolates belong to a distinct clade within the genus, denominated as clade I (Öztürk *et al.* 2020), and were the source of several linuron catabolic genes such as the linuron hydrolase-encoding *hylA* (Bers *et al.* 2013b) and *libA* (Bers *et al.* 2011a), the *dcaQTA1A2BR* cluster that encodes conversion of DCA to chlorocatechol, and the *ccdRCFDE* cluster encoding degradation of chlorocatechol to oxoadipate (Bers *et al.* 2011a; Bers *et al.* 2013b; Albers *et al.* 2017). This information was used to study the ecology of linuron biodegradation in agricultural soil. Using targeted PCR, *Variovorax* and its related linuron catabolic functions were shown to respond to linuron application in an agricultural field soil in Halen (Belgium), both in soil microcosms (MCs) and in the field. This soil was the origin of several of the above mentioned linuron-degrading *Variovorax* isolates and the ecological studies hence indicated that *Variovorax* effectively plays a role in the *in situ* dissipation of linuron (Bers *et al.* 2011b, 2012; Bers *et al.* 2013a; Breugelmans *et al.* 2007; Horemans *et al.* 2016; Sniegowski *et al.* 2011b; Sniegowski and Springael, 2015). However, other genera also harbour linuron-degrading soil isolates. Linuron-utilizing *Achromobacter* and *Hydrogenophaga* strains were isolated from the same Halen soil, of which the *Hydrogenophaga* strains depend on the same gene functions as *Variovorax* to degrade linuron, likely acquired by horizontal gene transfer (Breugelmans *et al.* 2007; Werner *et al.* 2020). Recent studies have further expanded the group of soil-borne linuron degraders with the isolation of a new consortium from soil near a pesticide producing factory in China. In this consortium, another β-proteobacterium, *Diaphorobacter*, performs the initial hydrolysis step through a novel hydrolase (Phh) (Zhang *et al.* 2018, 2020). A linuron transforming α-proteobacterium was recently isolated from herbicide-exposed soil and this *Sphingobium* strain uses yet another linuron hydrolase (LabB) (Zhang *et al.* 2018, 2020). As such, even though a plethora of information regarding the biodegradation of linuron exists, new degraders and gene functions involved in linuron degradation are still being identified, suggesting that the *in situ* ecology of linuron degrading bacteria in agricultural soil is more complex than currently known. Additionally, as mentioned above, potential biases associated with the enrichment and isolation of degraders from environmental samples, question the actual role of such linuron-degrading isolates in *in situ* biodegradation (Dealtry *et al.* 2016; Albers *et al.* 2017; Werner *et al.* 2020). This was highlighted by our recent study which suggested that cultivation might result in biased representation of *in situ* linuron degraders in an on-farm biopurification system (BPS) that treats pesticide contaminated wastewater. In that study, DNA-SIP identified Ramlibacter along with *Variovorax* as the main *in situ* assimilators in the examined BPS environment, while an EC obtained from the same BPS sample was dominated by *Variovorax* (Lerner *et al.* 2020).

This study revisits the above mentioned Halen soil questioning (i) whether *Variovorax* is indeed the main *in situ* linuron assimilator in this soil, (ii) whether cultivation and isolation led to biases in the identification of linuron degraders, and (iii) whether changes regarding the linuron-degrading community had occurred in the 5-10 year period between this study and earlier studies. To this end, new soil samples were collected from the Halen agricultural field and exposed to linuron in a MC set up similar to that used in previous studies, in order to stimulate growth of the linuron-degrading bacterial community. After 440 days of linuron dosing, the MC material was used for DNA-SIP combined with 16S rRNA gene amplicon sequencing and PCR analysis targeting known linuron catabolic genes and marker genes of mobile genetic elements (MGEs) that carry linuron catabolic genes. In addition, bacteria able to grow on linuron were isolated from an EC initiated from the same soil sample.

## Material and methods

### Chemicals used

Linuron (purity, 99.5%) was purchased from Sigma Aldrich (St. Louis, MO) and ^14^C linuron ([phenyl-U-^14^C]-linuron) (16.93 mCi mM^−1^, radiochemical purity >95%) from Izotop (Hungary). ^13^C linuron ([phenyl-U-^13^C] linuron) was synthesized as described before (Lerner *et al.* 2020).

### Microcosm setup

Soil was collected in November 2015 from an agricultural field in Halen, Belgium (50°55′40.2″N 5°06′59.3″E), that has been the subject of previous research (Bers *et al.* 2013a; Breugelmans *et al.* 2007; Sniegowski *et al.* 2011b). The field has been cultivated with rotating potato, wheat, and sugar beet crops throughout the years and has an unrecorded history of linuron treatment. Around 2 kg of top soil was collected at 2-20 cm depth from three locations in the field, approximately 100 m from one another (denominated as locations H1, H2 and H3), sieved (2 mm) and used to set up two sets of differently treated triplicate MCs per sample location, similar to those in previous studies (Sniegowski *et al.* 2011a; Lerner *et al.* 2020). The first set of MCs was supplemented three times per week with 1 mL of sterile Milli-Q^®^-water containing 20 mg L^−1^ linuron and was denominated H1^+^, H2^+^, and H3^+^ according to the field location. The second set was set up as a control to assess the impact of linuron addition and received sterile Milli-Q^®^-water without linuron (denominated as H1^−^, H2^−^, and H3^−^). All MCs were incubated at 25 °C in the dark. The linuron mineralization capacity in the MC materials was assessed after 9 months of treatment through a ^14^C-linuron mineralization assay as previously described (Sniegowski *et al.* 2011b).

### DNA-SIP analysis

DNA-SIP analysis was performed as described previously by Lerner *et al.* (2020) with material from the linuron supplemented MCs containing soil originating from location H3, taken after 440 days of treatment. Material was taken from the top 5 cm layer of each of the triplicate MCs, the three samples were pooled and mixed, and 0.5 g of the pooled material was weighed into seven 15-mL Pyrex tubes. Three of them subsequently received 0.5 mL of MS medium containing 100 mg L^−1^ of ^13^C-ring labeled linuron (^13^C-treatment), one received 0.5 mL of MS medium containing 100 mg L^−1^ of non-labeled (^12^C-) linuron (^12^C-treatment), and three received 0.5 mL of MS medium containing 100 mg L^−1^ of ^12^C-linuron spiked with 11.65 μg L^−1^ (~80 bq mL^−1^) ^14^C-linuron to determine linuron mineralization kinetics as reported before (Sniegowski *et al.* 2011b). After 7 days of incubation, DNA was extracted from the vials of the ^13^C- and ^12^C-treatment as reported before (Lerner *et al.* 2020). DNA extracts of two of the three ^13^C-treated replicates were pooled and the resulting pooled and non-pooled samples were further analyzed separately. Following isopycnic centrifugation and fractionation, DNA fractions relevant for 16S rRNA gene amplicon sequencing were identified by 16S rRNA gene based Denaturing Gradient Gel Electrophoresis (DGGE) community profiling. The density of the high density (heavy) DNA fractions selected for 16S rRNA gene amplicon sequencing fell within the previously reported range with an average density of ~ 1.72 g ml^−1^ (Dunford and Neufeld 2010; Pepe-Ranney *et al.* 2016). 16S rRNA gene amplicons were obtained, sequenced and the resulting data analyzed based on Amplicon Sequence Variants (ASVs) as described (Lerner *et al.* 2020). Only DNA fractions of the pooled ^13^C-treatment were sequenced since the 16S rRNA gene DGGE profiles obtained with the DNA fractions of pooled and un-pooled DNA were identical. DNA fractions of corresponding density of the ^12^C-treatment and non-fractionated DNA of both treatments were sequenced as well.

Genera involved in the assimilation of the linuron derived ^13^C, were identified by determining enrichment factors (EF) as described by Kramer *et al.* (2016) according to the formula

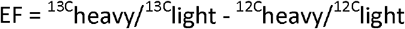

where ^13C^heavy and ^13C^light are the relative abundance of taxon-specific reads in heavy and light DNA fractions of ^13^C treatments, and ^12C^heavy and ^12C^light are the same for the respective ^12^C treatments. Only genera with a mean relative abundance of more than one percent across all samples were considered. Genera were considered as ^13^C labeled when the EF was higher than 0.5.

### Enrichment and isolation of linuron degraders

ECs growing on linuron as sole C-source, were initiated from the soil material collected at location H3 as described (Lerner *et al.* 2020), except that after the cultures were transferred twice, the cultures were plated on R2A plates. Collected single colonies were pre-grown in liquid R2B medium and the cells washed in MgSO_4_ and transferred, at a concentration of ~ 10^7^ cells mL^−1^, to 15-mL Pyrex tubes containing 5 mL MMO medium supplemented with 20 mg L^−1^ of linuron and 11.65 μg L^−1^ (~80 bq mL^−1^) ^14^C ring-labeled linuron, to monitor mineralization as described (Sniegowski *et al.* 2011b). Only strains that showed linuron mineralization were retained.

### PCR and DGGE analysis

The sequences of the primers used for PCR and the used PCR conditions are reported in Table S1 of Supplementary Material. PCR reactions were performed in a Biometra thermocycler (Analytik Jena, Germany) or Eppendorf Mastercycler (Eppendorf, Germany) apparatus. PCR products were visualized after agarose gel electrophoresis (1.5%) in Tris-acetate/EDTA buffer (90 V for 1 h) using GelRed. DGGE was performed in a C.B.S scientific apparatus (Del Mar, CA) as described (Moreels *et al.* 2004; Lerner *et al.* 2020). Pictures of DGGE profiles were edited using BioNumerics v7.6.3 (Applied Maths NV, St-Martens-Latem, Belgium).

### Sanger sequencing of 16S rRNA gene amplicons

16S rRNA gene amplicons were obtained from pure cultures of two isolates using primer set 27F and 1492R according to the described PCR conditions (Table S1) and sequenced on a Sanger platform using primer 1492R (GATC Biotech AG, Germany), recovering partial 16S rRNA gene sequences of around 1000 bps. Sequence similarity was determined using NCBI Blast (Morgulis *et al.* 2008). Maximum likelihood phylograms of relevant 16S rRNA gene sequences were obtained after alignment using Muscle v3.8.31 (Edgar 2004) and implementing a 1000x bootstrap analysis using RAxML v.8.2. (Stamatakis 2014).

### Nucleotide sequence accession numbers

The raw Illumina sequencing reads of the 16S rRNA amplicon sequencing and the 16S rRNA gene sequences of the obtained Isolates have been deposited under BioProject ID PRJNA657170 and GenBank accession numbers MT884016 and MT884017, respectively.

## Results

### Linuron mineralization in the Halen soil before and after treatment with linuron and selection of the DNA-SIP sampling point

Prior to linuron treatment, none of the soil samples from the three different sampling locations showed linuron mineralization (data not shown). However, supplementing the soils with linuron solution for 280 days, resulted in a clear linuron mineralizing capacity with a mineralization efficiency between 30% and 46% in the ^14^C-linuron mineralization assay. In contrast, no linuron mineralization was recorded in the MCs that received linuron-free MilliQ-water (Figure S1). The linuron treated MCs incubated with soil from location H3 (MC set H3^+^) showed the highest mineralization capacity and was used for DNA-SIP analysis after 440 days of linuron treatment. To verify linuron mineralization and to determine linuron mineralization progress in order to select the appropriate time point for DNA-extraction, a ^14^C-mineralization assay was performed in paralell. ^14^CO_2_ production stabilized close to 40%, indicating clear mineralization of the added linuron and the likely incorporation of residual carbon into linuron mineralizing biomass. In addition, the observed mineralization kinetics fit the Gompertz kinetic model, suggesting that the compound was used for growth (Figure 1A) (Johnsen *et al.* 2013). Extended incubation times can increase decay of primary degraders through apoptosis or predation, leading to cross-labeling of secondary populations, while short incubation times can result in insufficient labeling of primary degaders for detection (Neufeld *et al.* 2007). We decided to perform the DNA extraction for DNA-SIP analysis on samples taken at day 7, when mineralization, and likely the growth of linuron assimilators, reached the late exponential phase, when maximum linuron mineralization was realized and sufficient growth of primary degraders with minimal decay was expected.

**Figure 1.**
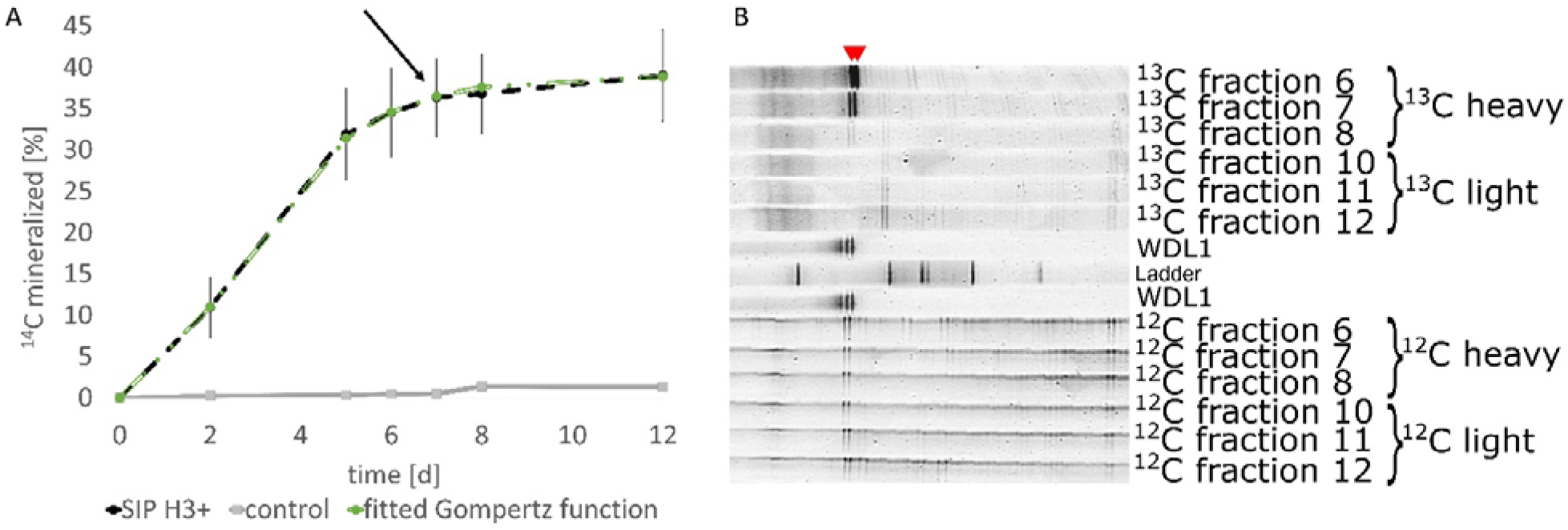
**A)** ^14^C-linuron mineralization kinetics observed in samples taken at day 440 of linuron dosing from the MCs containing soil from location H3 performed in parallel to the ^13^C-treatment for DNA-SIP analysis. The kinetics of ^14^C-linuron mineralization was fitted to the Gompertz function (green line). The arrow indicates the time point of DNA extraction, i.e. 7 days after adding ^13^C-linuron, for further SIP analysis. The control contains medium without addition of soil. ^14^C mineralization refers to the % of produced ^14^CO_2_ in reference to the initial amount of added ^14^C-linuron. The error bars indicate the standard deviations of three replicates. **B)** 16S rRNA gene amplicon DGGE showing community fingerprints recovered from the CsCl-separated DNA fractions of the ^13^C- and ^12^C-treatments of the DNA-SIP. Results are shown for the pooled DNA of two replicate DNA-SIP assays. The top of the gel shows six fractions of the ^12^C-treatment, while the bottom of the gel shows six fractions of the ^12^C-treatment. In the center, an in-house reference ladder and two lanes showing DGGE profiles obtained from DNA of the linuron degrading isolate *Variovorax* WDL1 (WDL1) are shown. The positions of the bands that correspond to WDL1 are marked by red arrows. The fractions determined as heavy and light, which were pooled and used for 16S rRNA gene amplicon sequencing analysis, are indicated accordingly. Only DNA fractions that were sequenced are shown.

### Analysis of relevant DNA-SIP density fractions

The DNA extracts of the ^13^C- and ^12^C-treatment were subjected to isopycnic centrifugation and the collected fractions analyzed by bacterial 16S rRNA gene amplicon DGGE. Results of the pooled DNA (of two replicate DNA-SIP assays) are shown in Figure 1B. Heavy DNA fractions of the ^13^C-treatment exhibited a prominent banding pattern, showing distinct DGGE bands that migrated similarly to those obtained with DNA of the linuron degrading *Variovorax* strain WDL1. These bands were absent in the light fractions, indicating enrichment of specific organisms assimilating the added ^13^C-linuron. The DNA fractions of the ^12^C-treatment showed the same distinct banding pattern, but the bands were less intense and evenly spread across the density gradient, suggesting that the occurrence of these bands in the heavy fractions of the ^13^C-treatment is due to assimilation of ^13^C-linuron rather than to the existence of naturally high DNA density in the corresponding organisms (Figure 1B). Identical results were obtained with the non-pooled DNAs (Figure S2). Three heavy fractions of the ^13^C-treatment exhibiting the distinct banding patterns and three light fractions were selected for further 16S rRNA gene amplicon sequencing, as well as the corresponding fractions of the ^12^C-treatment (Figure 1B). In all cases, the respective selected fractions of similar density were pooled prior to amplicon sequencing, resulting in one heavy and one light fraction of both the ^13^C- and ^12^C-treatment. In addition, non-fractionated DNA of both the ^13^C- and ^12^C-treatment was analyzed.

ASV analysis revealed a high similarity between all samples regarding community composition, except for two ASV groups that were clearly enriched in the heavy DNA fractions of the ^13^C-treatment (Figure 2). Both were identified as *Variovorax* (designated as *Variovorax* I and *Variovorax* II) and represented, respectively, 4% and 2% of the total sequence reads in the heavy DNA fractions of the ^13^C-treatment. The ASVs were enriched by a factor of 190 and 31 compared to the light fractions, and by a factor of 10 and 18 compared to the non-fractionated DNA. Furthermore, they were 7- and 20-fold more abundant in the heavy ^13^C treated fractions than in the heavy ^12^C treated fractions. Calculated EFs for the two *Variovorax* ASVs were ~190 and ~30 respectively, confirming their designation as ^13^C-labeled and hence as linuron assimilating organisms (Figure 2). The sequenced 16S rRNA gene region of *Variovorax* I was identical to those of *Variovorax* clade I organisms that harbor all known linuron degrading *Variovorax* isolates such as *Variovorax* sp. WDL1, PBS-H4, and PBL-H6 (Öztürk *et al.* 2020), of which the latter two strains were previously isolated from a Halen soil sample taken in 2005 (Breugelmans *et al.* 2007). This ASV was furthermore identical to cloned *Variovorax* 16S rRNA gene sequences obtained from a culture enriched on linuron initiated with BPS material (Lerner *et al.* 2020). The *Variovorax* II sequence did not match any known linuron degrader, but was instead identical to the V4 16S rRNA gene region of the DCA degrading isolate *Variovorax* sp. PBD-E37 recovered from an EC derived from a Danish agricultural soil and belonging to *Variovorax* clade II that contains most of the *Variovorax* species identified up to now but no linuron degrader (Breugelmans *et al.* 2007; Öztürk *et al.* 2020).

**Figure 2.**
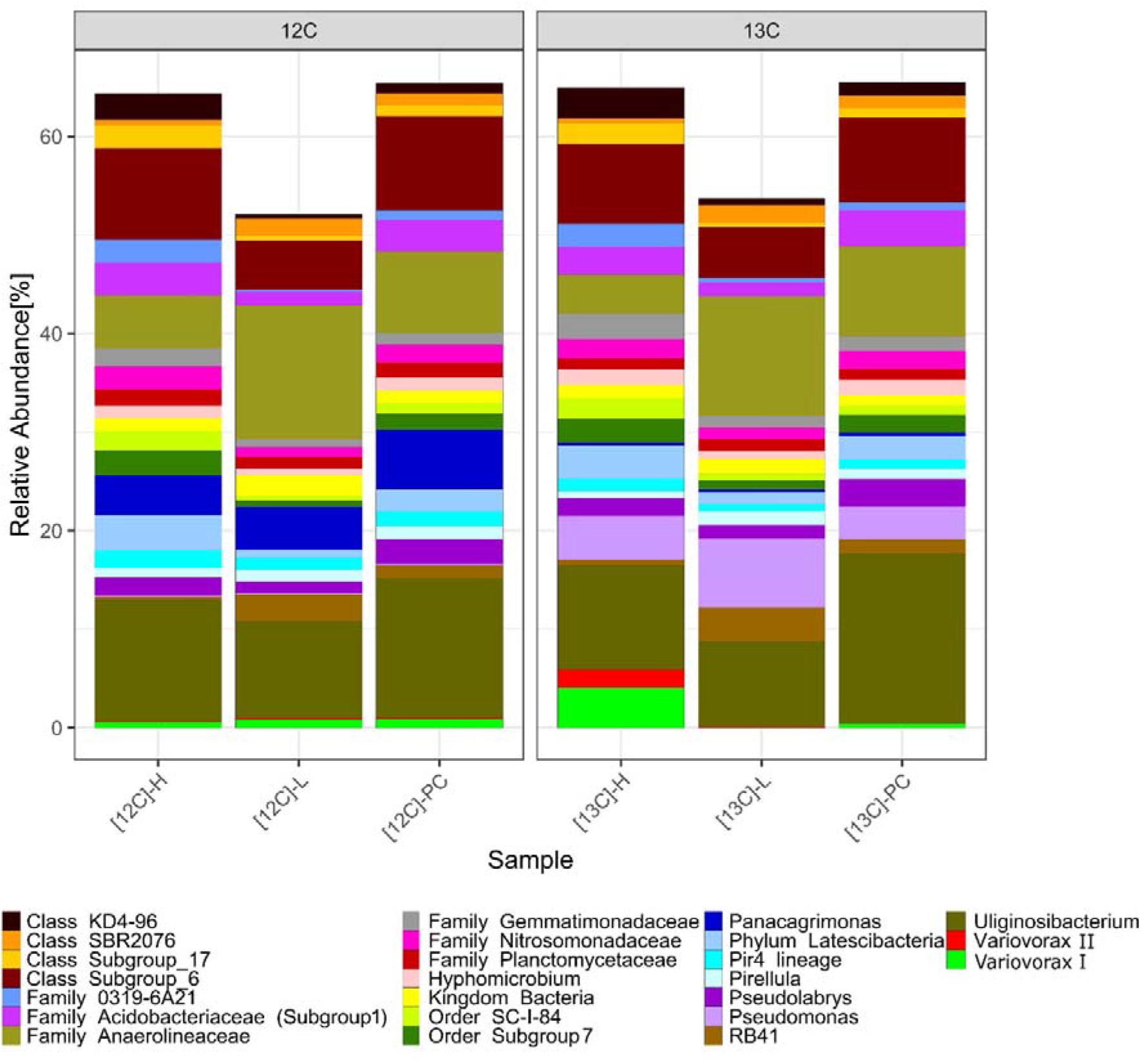
Relative abundance of ASVs and their taxonomic classification at the genus level in the heavy (H) and light (L) DNA density fractions obtained from either ^12^C-treatment (12C) or ^13^C-treatment (13C) samples, as well as of non-fractionated DNA samples (PC). Only ASVs with a mean abundance of >1% across all samples or >1% abundance in the heavy ^13^C DNA-SIP sample were considered. ASVs of the same genus were pooled for visualization, with exception of the two *Variovorax* ASVs indicated as *Variovorax* I and *Variovorax* II.

### PCR detection of marker genes for linuron degradation and horizontal gene transfer in DNA-SIP fractions

PCR amplification was applied to the different DNA fractions to detect (i) genes that encode essential steps in the linuron degradation pathway, i.e. linuron hydrolysis genes *hylA* and *libA* and two variants of the *dcaQ* gene (*dcaQI* and *dcaQII*) involved in the conversion of the hydrolysis product DCA to chlorocatechol, and (ii) marker genes for MGEs known to carry or to be associated with the aforementioned catabolic genes, i.e. *trfA*, as a marker of IncP-1 plasmids and *tnpA* as a marker for the insertion sequence IS1071 (Albers *et al.* 2017; Dealtry *et al.* 2016; Dunon *et al.* 2018). A clear enrichment of *hylA* and the *dcaQ* gene variant *dcaQII*, both first identified in the linuron degrading *Variovorax* sp. WDL1, was found in the heavy DNA fractions of the ^13^C-treatment, while their amplifaction in the ^12^C-gradient occured mainly in the lighter fractions (Figure 3). For *libA* only aspecific amplification was observed, while the dcaQ gene variant *dcaQI* that exists in conjunction with *libA* in the linuron degrading *Variovorax* sp. SRS16, was enriched in fractions 7-9 of the ^13^C-treatment. In the ^12^C-treatment, *dcaQI* was only amplified in the light fractions. The MGE marker genes *trfA* and *tnpA* showed strong signals along both the ^12^C and ^13^C DNA-SIP gradients, but a clear enrichment of *tnpA* was observed in the heavy fractions of the ^13^C-treatment while this was not the case in the ^12^C-treatment (Figure 3).

**Figure 3.**
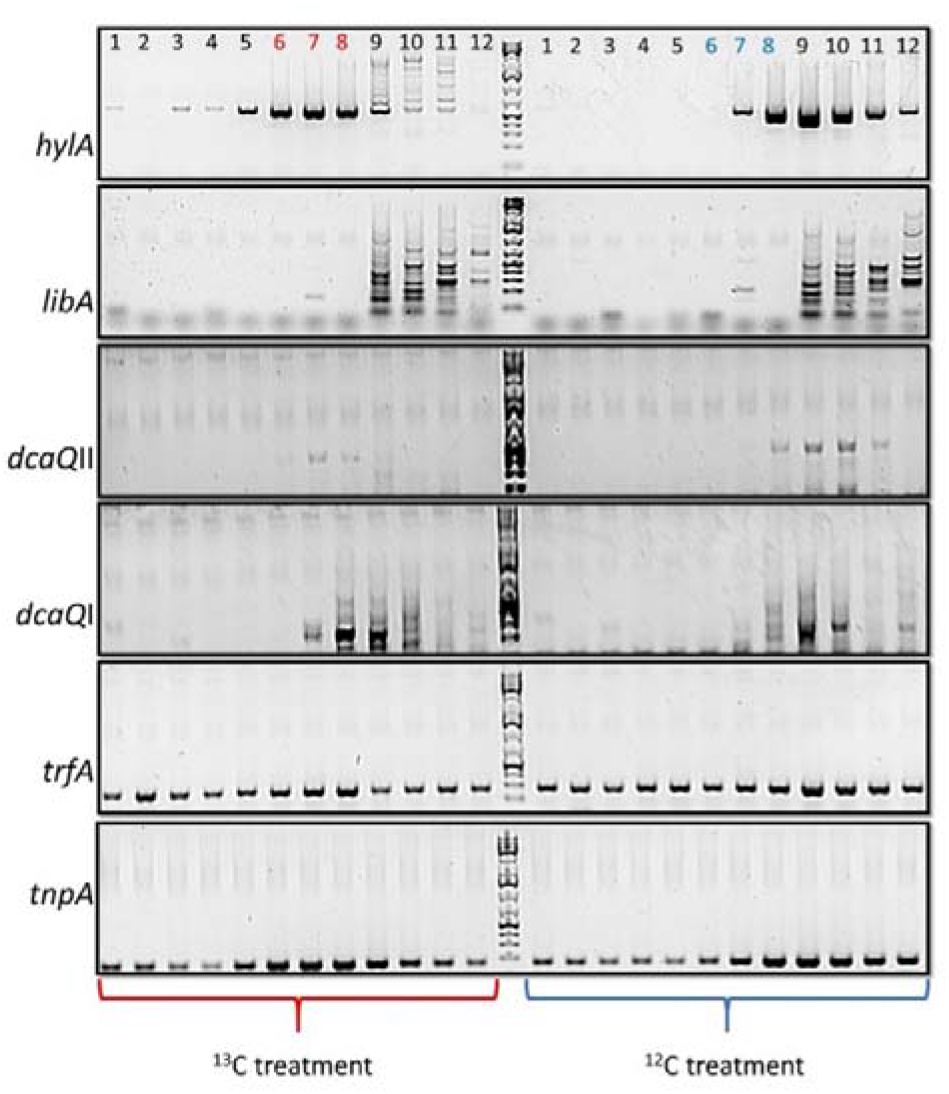
PCR analysis targeting linuron catabolic and MGE marker genes in the DNA-SIP fractions of the ^13^C- and ^12^C-treatment. The DNA fractions of the ^13^C-treatment are marked in red below the gel and those of the ^12^C-treatment in blue. The numbers above the lanes indicate the DNA fractions with density decreasing from 1-12. The lane labels indicated in red and blue mark the heavy fractions selected for 16S rRNA gene amplicon sequencing from the ^13^C- and ^12^C-treatment, respectively.

### Isolation of linuron mineralizing strains from Halen agricultural soil

An EC that rapidly degraded linuron was obtained from the Halen soil of location H3. From the EC, two bacterial isolates were recovered which mineralized linuron in pure culture (Figure S3). The isolates were tested for the presence of *hylA* and *libA* by PCR and tested positive for *libA*. Sanger sequencing of the 16S rRNA gene, yielding an approximately 1000-bp long portion of the gene, identified both isolates as *Variovorax* with a close similarity (>99.5%) to linuron degrading isolates such as *Variovorax sp*. WDL1, PBS-H4, and PBL-H6. Phylogram calculation positioned the isolated *Variovorax* strains within the previously described linuron degrading *Variovorax* clade I (Figure 4) (Öztürk *et al.* 2020). The sequences further exhibited a close similarity to *Variovorax* 16S rRNA sequences retrieved through cloning from a linuron degrading culture enriched from BPS material (>99.3%) and were 100% identical to the V4 region of the 16S rRNA gene of the *Variovorax* I ASV, appointed as linuron assimilating organism in the DNA-SIP (Figure 2).

**Figure 4.**
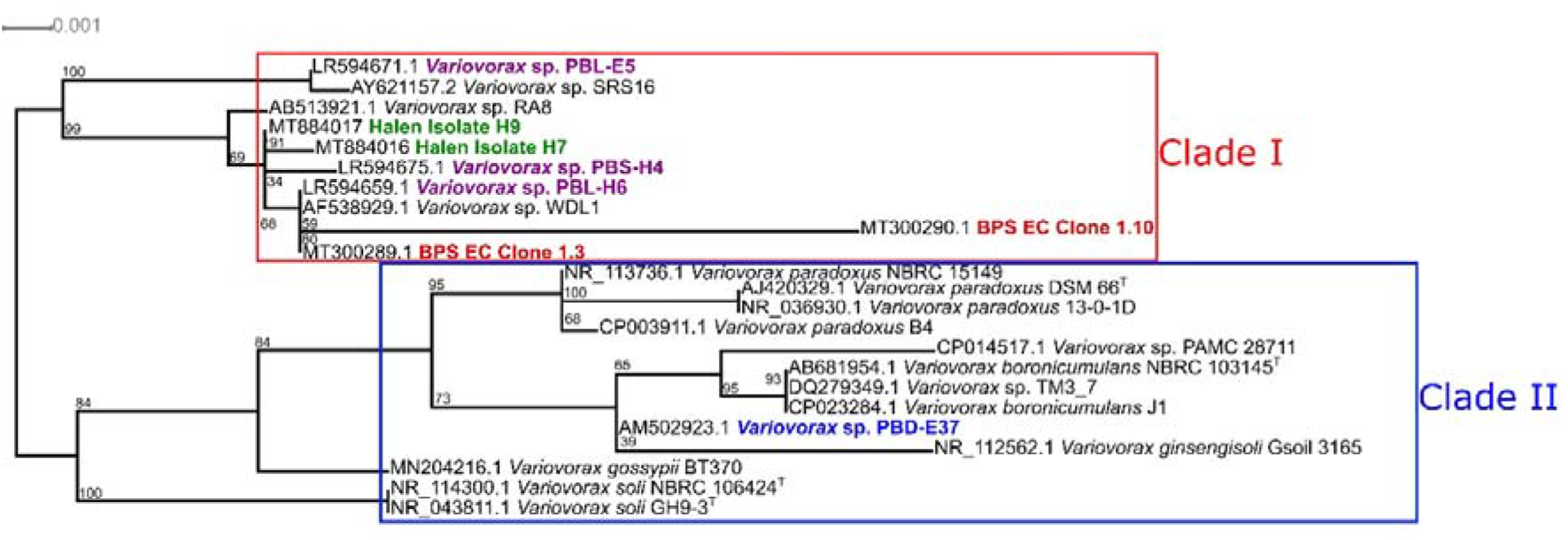
Maximum likelihood phylogram based on ~1kb partial 16S rRNA gene sequences (bp 300 to bp 1300) positioning the linuron degrading *Variovorax* isolates H9 and H7, obtained from the Halen soil (green), within the *Variovorax* genus. Linuron-degrading *Variovorax* sp. strains isolated from the Halen soil in previous studies are indicated in purple, while two cloned partial *Variovorax* 16S rRNA gene sequences (EC clones 1.3 and 1.10) obtained from a linuron degrading EC initiated from BPS material are marked in red and the DCA degrading *Variovorax* sp. PBD-E37 is marked blue. Branches corresponding to the reported *Variovorax* clades I and II are indicated in red and blue, respectively.

## Discussion

### DNA-SIP identifies Variovorax strains as in situ assimilators of linuron in the Halen agricultural soil

Two *Variovorax* ASVs, *Variovorax* I and *Variovorax* II, were revealed as the main *in situ* assimilators of linuron in the Halen agricultural soil. Previous studies on soil samples from the same agricultural field, collected 10 and 5 years earlier, also appointed *Variovorax* as a key player in linuron removal. Using soil collected in 2005, Breugelmans *et al.* (2007) isolated several *Variovorax* strains that were able to degrade linuron, DCA, or both. Bers *et al.* (2011b) provided further evidence of the importance of *Variovorax* in linuron removal in the same soil sample through cultivation free analysis via targeted (q-)PCR and DGGE analysis, showing that the *Variovorax* community positively responded to linuron application in a MC setup. The results were confirmed and expanded in a field study in 2010 and a concomitant MC study using soil taken at the start of the field study (Bers *et al.* 2013a; Horemans *et al.* 2016). While these studies suggested a clear link between linuron dissipation capacity and *Variovorax*, the used approaches only provide evidence regarding their presence and response but do not unambiguously inform about their direct participation in *in situ* degradation. Moreover, other organisms besides *Variovorax* might be involved in linuron degradation and their response would be missed by the targeted PCR approach. For instance, Dealtry *et al.* (2016) studied the response of a BPS community to linuron application and found that apart from the *Comamonadaceae* family, to which *Variovorax* belongs, several other families, such as Hyphomicrobiaceae, Sphingomonadaceae and *MethylophylAceae* increased in relative abundance. Similarly, Lerner *et al.* (2020) using DNA-SIP, revealed the involvement of *Ramlibacter* and an unclassified Comamonadaceae genus in linuron dissipation, next to *Variovorax*, in a BPS community. The current study for the first time unambiguously demonstrates that *Variovorax* is the main genus assimilating the linuron-derived ^13^C *in situ* in the Halen agricultural field. Both *Variovorax* ASVs identified by DNA-SIP, showed high similarities to previously described *Variovorax* isolates from linuron exposed agricultural soils. The sequence of *Variovorax* I was identical to that of the linuron-degrading *Variovorax* isolates PBS-H4 and PBL-H6, previously isolated from the Halen soil (Breugelmans *et al.* 2007), as well as to *Variovorax* sp. WDL1 isolated from a linuron exposed soil in Gorsem (Belgium) and two cloned 16S rRNA gene sequences obtained from a linuron degrading EC initiated with BPS material (Dejonghe *et al.* 2003; Lerner *et al.* 2020). All these strains belong to the distinct *Variovorax* clade I that harbours all linuron hydrolysing *Variovorax* isolates described so far and hence seems to be prone to adapting towards linuron catabolism (Öztürk *et al.* 2020). In contrast, the sequenced 16S rRNA gene region of *Variovorax* II matched that of the DCA-degrading isolate *Variovorax* sp. PBD-E37, isolated from a Danish agricultural soil, which belongs to *Variovorax* clade II, harbouring the majority of isolated *Variovorax* species including *Variovorax* paradoxus. Several other genera than *Variovorax*, i.e. *Hydrogenophaga*, *Achromobacter*, *Diaphorobacter*, *Sphingobium*, Mycobacterium and Arthrobacter have been isolated as linuron degraders in either the investigated soil or in other environments. However, none of them were identified as *in situ* linuron assimilators using the DNA-SIP analysis. As such, this study underlines the importance of *Variovorax* in the bacterial ecology of linuron removal in agricultural soil and particularly the Halen soil. Moreover, assuming that *Variovorax* was the main *in situ* linuron-degrader in the samples of 2005 and 2010, our study highlights the robustness of *Variovorax* as a linuron degrader in the Halen soil and shows that, once established, xenobiotic catabolic strains and gene functions can be maintained and memorized in a soil community for an extended period of time, including periods without linuron application. Studies on the long-term robustness of established pesticide degrading populations and gene functions are rare. One three year field study suggested a similar memory effect, showing the long-term survival of an atrazine-degrading bacterial population in an atrazine-treated soil, despite periods without atrazine application, and their proliferation once the compound was re-applied (Cheyns *et al.* 2012).

### Linuron assimilation in the Halen agricultural soil is linked to known linuron catabolic genes and MGEs and likely involves a synergistic interaction

Targeted PCR revealed the involvement of linuron catabolic genes *hylA*, *dcaQI* and *dcaQII* in the *in situ* degradation of linuron, as indicated by their enrichment in the heavy fractions of the ^13^C-treatment. Interestingly, the alternative *Variovorax* associated linuron hydrolase gene *libA* could not be linked to *in situ* linuron degradation, despite being present as shown by the isolation of *libA*-containing linuron degraders from the soil. The enrichment of *hylA* over *libA* in the current study is in line with previous studies where *hylA* was 100 to 1000-fold more abundant in the Halen soil following linuron application, both in soil MCs and in field studies (Horemans *et al.* 2016). Similar results were recently reported in a MC based DNA-SIP study of a BPS environment, where initially both *libA* and *hylA* were detected, but after longer linuron dosing, only *hylA* remained detectable (Lerner *et al.* 2020). Apparently, linuron degraders associated with *hylA* have an increased fitness in the Halen soil and other environments like the BPS environment. *HylA* and *LibA* are isofunctional linuron hydrolases that belong to different hydrolase families and differ in biochemical characteristics, but whether and how these differences govern the fitness of their hosts is currently unknown (Bers *et al.* 2013b). Similarly, *dcaQI* and *dcaQII* are variants of a gene that represents two isofunctional dca clusters, respectively. The two variants exhibit substantial amino acid sequence dissimilarities ranging from 16 to 30% in the proteins composing the DCA dioxygenase and differences might be expected in biochemical properties, although there is currently no experimental evidence available supporting that hypothesis. In an earlier study *dcaQI* and *dcaQII* were also reported to respond to linuron application in the Halen soil (Horemans *et al.* 2016), but *dcaQII* became more abundant than *dcaQI*, while in this study the intensity of the PCR signals indicates the opposite. Moreover, our recent analysis of the genome sequences of linuron-degrading *Variovorax* strains show that *dcaQII* is also present as a truncated version just upstream of *hylA* (Albers *et al.* 2017; Öztürk *et al.* 2020). This truncated version includes binding sites for the *dcaQII* primers used in this study for PCR and hence we cannot exclude that the observed PCR signal might stem from a non-functional truncated *dcaQII* gene that becomes enriched due to its linkage to *hylA*. The recently reported catabolic genes like phh and labB were not tested in this study and can therefore not be excluded from being involved in *in situ* biodegradation in the Halen soil (Zhang *et al.* 2018, 2020).

Interestingly, to date, all linuron-degrading isolates that carry *hylA*, like strains PBS-H4 and WDL1, do not carry a full dca cluster. Strain WDL1 was originally reported to carry a full *dcaQII* containing dca gene cluster but the strain was recently shown to consist of two subpopulations, one carrying *hylA* and one carrying the dca gene cluster (Dejonghe *et al.* 2003; Albers *et al.* 2017). Moreover, strains like WDL1 and PBS-H4 were always recovered as members of consortia. In these consortia they perform the crucial linuron hydrolysis step, but depend on additional strains for further linuron mineralization, including the dca encoded DCA dioxygenase step. The latter belong to genera like Comamonas, *Variovorax* and *Delftia* and can contain either one of the two dca variants (Öztürk *et al.* 2020). The observed enrichment of *hylA* and dca in the DNA-SIP might therefore indicate that *in situ* linuron catabolism in the Halen soil involves a similar consortium. The enrichment of two *Variovorax* ASVs in the DNA-SIP analysis supports this. While *Variovorax* I, which is most related to strains WDL1 and PBS-H4, would perform the conversion of linuron to DCA, *Variovorax* II would then provide the dca dependent DCA conversion step. The latter is further supported by the close sequence similarity of *Variovorax* II to *Variovorax* sp. PBD-E37, a DCA degrader previously obtained from a linuron degrading consortium. Moreover, the 16S rRNA genes of *Variovorax* sp. PBD-E37 and *Variovorax* II cluster within the *Variovorax* clade II and therefore the species corresponding to *Variovorax* II is unlikely to carry the *hylA* gene, strengthening the hypothesis that the role of *Variovorax* II in *in situ* linuron degradation in the Halen soil rather lies in downstream processing of linuron hydrolysis intermediates. However, the question where the organism represented by the *Variovorax* II ASV acquires its assimilated ^13^C from remains. The linuron used for DNA-SIP is ^13^C-labelled in the ring structure and organisms that hydrolyze linuron but lack the dca cluster do not necessarily assimilate the ^13^C unless they retrieve it from a DCA degrading organism. This is reminiscent of the observations made in the linuron degrading consortium from which strain WDL1 originated. In that consortium, the subpopulation of WDL1 that only contains *hylA* and lacks the dca cluster, shows clear growth when linuron is supplied as the sole C-source and hence receives a substantial amount of the carbon of the aromatic ring. A plausible explanation is that part of the chlorocatechol resulting from the conversion of DCA is utilized by WDL1 since it contains the ccd gene cluster that converts chlorocatechol to Krebs cycle intermediates. Such a scenario would also explain the ^13^C assimilation by both *Variovorax* populations in the DNA-SIP.

In all linuron-degrading isolates, the examined catabolic genes are located on plasmids of high promiscuity, like IncP-1 and PromA, or on larger megaplasmids with unknown promiscuity (Dealtry *et al.* 2016; Öztürk *et al.* 2020; Werner *et al.* 2020). On these plasmids, the linuron catabolic genes, either encoding the hydrolase, DCA dioxygenase or chlorocatechol catabolism, are always embedded in a composite transposon characterized by flanking IS1071 insertion sequences (Dunon *et al.* 2013, 2018b). The enrichment of its marker gene *tnpA* in the heavy ^13^C fractions that also showed enrichment of *hylA*, *dcaQI*, *dcaQII*, and *Variovorax* was evident, suggesting that at least some of the catabolic genes in the active *in situ Variovorax* populations are associated with IS1071. This agrees with a recent metagenomics study that physically linked the different linuron catabolic genes to IS1071 in the Halen soil (Dunon *et al.* 2018b). In a BPS environment the linuron catabolic genes were also linked to IS1071, both based on metagenomics and DNA-SIP (Dunon *et al.* 2018b; Lerner *et al.* 2020). Therefore, the linkage with *IS1071* appears a ubiquitous characteristic of the *Variovorax* linuron catabolic genes. The results for IncP-1 were less clear, but the catabolic genes might have been associated with other (conjugative) plasmids, as is the case in *Variovorax* sp. PBS-H4 that hosts a PromA plasmid carrying *hylA* and the *ccd* cluster and in the strain WDL1 populations that carry a megaplasmid containing *hylA* or the dca cluster, as well as the *ccd* cluster (Breugelmans *et al.* 2007; Öztürk *et al.* 2020; Werner *et al.* 2020).

### Soil free enrichment of linuron-degrading cultures yields *libA*-carrying *Variovorax* isolates but not *hylA*-containing strains

Two isolates that mineralize linuron were obtained from an EC initiated with the same soil sample that was used for the MC set up and DNA-SIP. Based on their 16S rRNA gene sequences, the isolates were related to previously reported linuron-degrading *Variovorax* isolates such as WDL1, PBS-H4, and PBL-H6 (>99.6%) (Dejonghe *et al.* 2003; Breugelmans *et al.* 2007), of which PBS-H4 and PBL-H6 originated from the same agricultural field. Remarkably, both isolates obtained in the current study carried *libA* instead of *hylA*. This observation might indicate that *libA* containing *Variovorax* were privileged in the soil free EC rather than the *hylA* containing *Variovorax* and that soil free enrichment results in a biased representation of the *in situ* linuron-assimilating bacterial populations. The isolation procedure was, however, directed towards the isolation of linuron mineralizing bacteria, hence organisms that perform the complete pathway and, as mentioned above, up to date this only applies to *libA*-containing *Variovorax* strains. As such, *hylA*-containing *Variovorax*, despite being present in the EC, might have been excluded from isolation because they are unable to mineralize linuron. Curiously, the *libA* containing strain PBL-H6 was isolated from 2005 samples using a similar soil free enrichment procedure as used in the current study, while the *hylA* containing PBS-H4 was isolated as part of a consortium using a slightly different procedure in which soil particles were retained in the EC (Breugelmans *et al.* 2007). On the other hand, soil free linuron-degrading enrichments initiated from 2010 samples were dominated by *hylA*-carrying *Variovorax* strains instead of *libA*-containing *Variovorax* strains (Bers *et al.* 2013a). Likely, subtle differences in enrichment conditions steer the community to either *libA*- or *hylA*-containing microbes. Nevertheless, the finding that highly similar *libA*-containing *Variovorax* strains were isolated at two different time points a decade apart, emphasizes the robustness of these organisms and the existence of a strong genetic memory regarding linuron biodegradation in the Halen soil.

## Conclusion

The revisiting of the Halen agricultural soil for the first time conclusively identified *Variovorax* as the main *in situ* assimilator of linuron-derived carbon in an agricultural soil environment. Full linuron degradation is likely achieved due to a collaboration between two *Variovorax* species that exchange linuron-derived carbon for growth. Moreover, the study strongly suggests the involvement of *hylA* and the dca cluster, likely encompassed in a composite transposable element, in *in situ* biodegradation. As previously observed, soil free enrichment and isolation of linuron mineralizing strains from the same soil sample resulted in the isolation of similar *Variovorax* strains containing *libA* rather than *hylA*, indicating a possible bias of soil free enrichment procedures in identifying *in situ* linuron degraders. Overall, the study points to the existence of a strong genetic memory regarding linuron degradation that is linked to *Variovorax* in the examined Halen soil and was retained for more than a decade, despite periods with complete absence of linuron application.

## ACKNOWLEDGEMENTS

The authors thank S. Bettens for collection of the soil samples and set up of the MCs, B. Müller for assistance in conducting DNA-SIP and D. Grauwels for overall technical assistance. This work was supported by the FWO [Project G.0371.06], the EU project METAEXPLORE [EU grant n°222625] and the KU Leuven [Project C14/20/063].

