## Supplementary data for "DNA-SIP and repeated isolation corroborate *Variovorax* as a key organism in maintaining the genetic memory for linuron biodegradation in an agricultural soil"

**Table S1.** PCR conditions and primers used in this study

| Target gene / primer | Primer sequence [5’ – 3’] | Program | | | | | | | | | | | | | | |
| --- | --- | --- | --- | --- | --- | --- | --- | --- | --- | --- | --- | --- | --- | --- | --- | --- |
|  |  | Initial denaturation | | Denaturation | | | Annealing | | | | Elongation | | | | Final elongation | |
| *hylA* (Lerner *et al.* 2020) | GCAGGGGCTAACAGTGAAGT | 95°C | 5m | 95°C | 30s | | 55°C | | 30s | | 72°C | | | 1m | 72°C | 10m |
|  | TGAAGGTCATGTCCACTCGC |  |  | 30x | | | | | | | | | | |  |  |
| *libA* (Lerner *et al.* 2020) | GTTCATTCCTCCGGCAGACA | 95°C | 5m | 95°C | 30s | | 55°C | | 30s | | 72°C | | | 1m | 72°C | 10m |
|  | GCCGTGAAAGGAATCGCATC |  |  | 30x | | | | | | | | | | |  |  |
| *dcaQI* (Horemans *et al.* 2016) | CTCTCATGGCCGGATCAATA | 95°C | 5m | 95°C | 30s | | 55°C | | 30s | | 72°C | | | 1m | 72°C | 10m |
|  | TACAGATCGGCCAGCATCCA |  |  | 30x | | | | | | | | | | |  |  |
| *dcaQII* (Horemans *et al.* 2016) | CGCCCACTGGTCATGTAAAG | 95°C | 5m | 95°C | 30s | | 55°C | | 30s | | 72°C | | | 1m | 72°C | 10m |
|  | GAAAAGCACGGCATCTGGTC |  |  | 30x | | | | | | | | | | |  |  |
| *trfA* (α,β,ε) (Bahl *et al.* 2009) | TTCACSTTCTACGAGMTKTGCCAGGAC | 95°C | 5m | 94°C | 30s | | 60°C | | 20s | | 72°C | | | 20s | 72°C | 5m |
|  | GWCAGCTTGCGGTACTTCTCCCA |  |  | 35x | | | | | | | | | | |  |  |
| *tnpA* (Providenti *et al.* 2006) | GCTTGGTCACTTCTGGGTCTTC | 95°C | 3m | 95°C | 35s | | 58°C | | 35s | | 72°C | | | 1m | 72°C | 10m |
|  | CTATGCCCGTCTATCGTTACCC |  |  | 35x | | | | | | | | | | |  |  |
| 16S rRNA gene 27F / 1492R (Turner *et al.* 1999) | AGAGTTTGATCCTGGCTCAG | 95°C | 5m | 95°C | 1m | | 52°C | | 1m | | 72°C | | | 1.5m | 72°C | 8m |
|  | TACGGYTACCTTGTTACGACTT |  |  | 35x | | | | | | | | | | |  |  |
| 16S rRNA gene 63F / 518R (El Fantroussi *et al.* 1999) | CAGGCCTAACACATGCAAGT | 95°C | 3m | 95°C | 30s | | 58°C | | 30s | | 72°C | | | 45s | 72°C | 5m |
|  | ATTACCGCGGCTGCTGG |  |  | 35x | | | | | | | | | | |  |  |
| 16S rRNA gene 8F / 533R (Klindworth *et al.* 2013) | AGAGTTTGATYMTGGCTCAG | 95°C | 3m | 95°C | 30s | | 58°C | | 30s | | 72°C | | | 45s | 72°C | 5m |
|  | TTACCGCGGCTGCTGGCAC |  |  | 35x | | | | | | | | | | |  |  |
| 16S rRNA gene GM3F / 908R(Klindworth *et al.* 2013) | AGAGTTTGATCMTGGC | 95°C | 3m | 95°C | 30s | | 50°C | | 30s | | 72°C | | | 1m | 72°C | 10m |
|  | CGTCAATTCMTTTGAGTT |  |  | 30x | | | | | | | | | | |  |  |
| 16S rRNA gene A519F / 802R(Klindworth *et al.* 2013) | CAGCMGCCGCGGTAA | 95°C | 3m | 95°C | | 30s | | 55°C | | 30s | | 72°C | 30s | | 72°C | 5m |
|  | TACNVGGGTATCTAATCC |  |  | 25x | | | | | | | | | | |  |  |

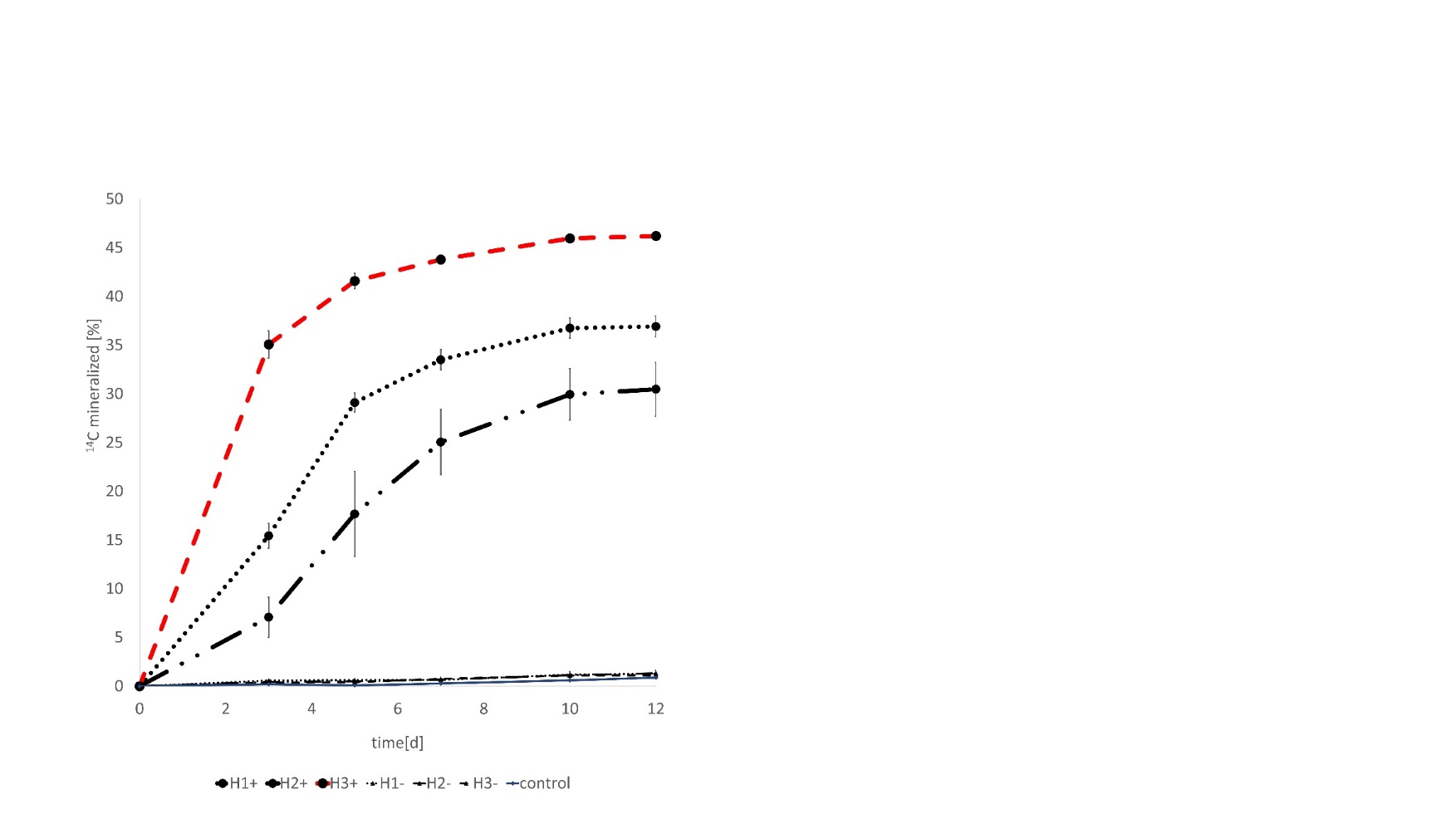

**Figure S1.** ^14^C-linuron mineralization kinetics in samples taken from the MCs containing Halen soil after 280 days of treatment with linuron-supplemented MilliQ water (+) or with linuron-free MilliQ water (-). Soil was sampled from three different locations within the Halen field H1, H2 and H3. The control contains medium without soil. ^14^C mineralization in the Y-axis refers to the % of produced ^14^CO_2_ in reference to the initial amount of added ^14^C-linuron. Averages of three replicates are shown with the error bars indicating the standard deviations.

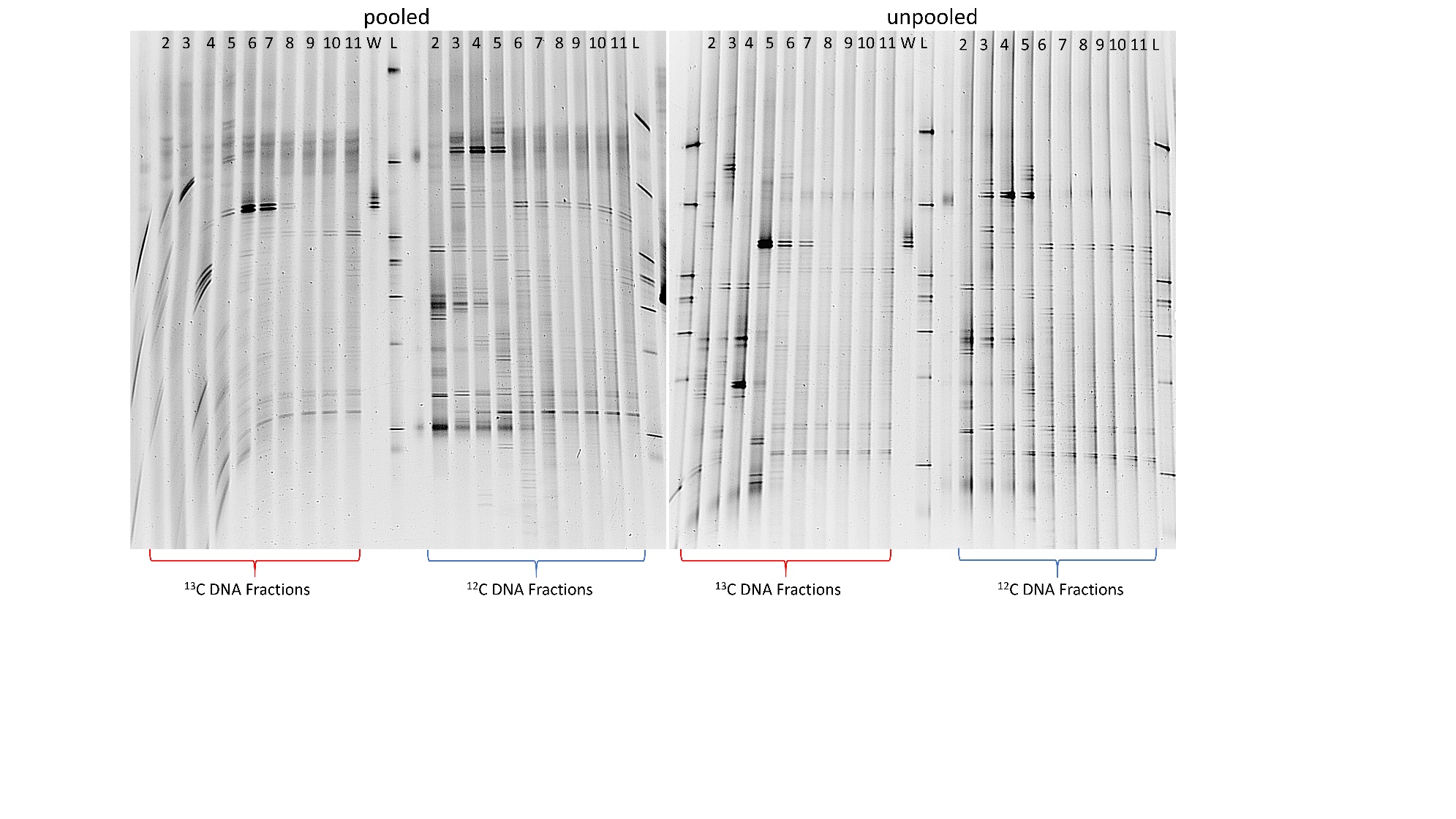

**Figure S2.** 16S rRNA gene amplicon DGGE showing community fingerprints recovered from the CsCl-separated DNA fractions of the DNA-SIP using the non-pooled DNA of one replicate. The ^13^C- and ^12^C-treatments are indicated in red and blue below the gel. Ten fractions (2-11) of the ^13^C- and ^12^C-treatment are depicted. The DGGE band obtained with DNA of *Variovorax* sp. WDL is labeled as “W” while the in-house ladder is labelled as “L”.

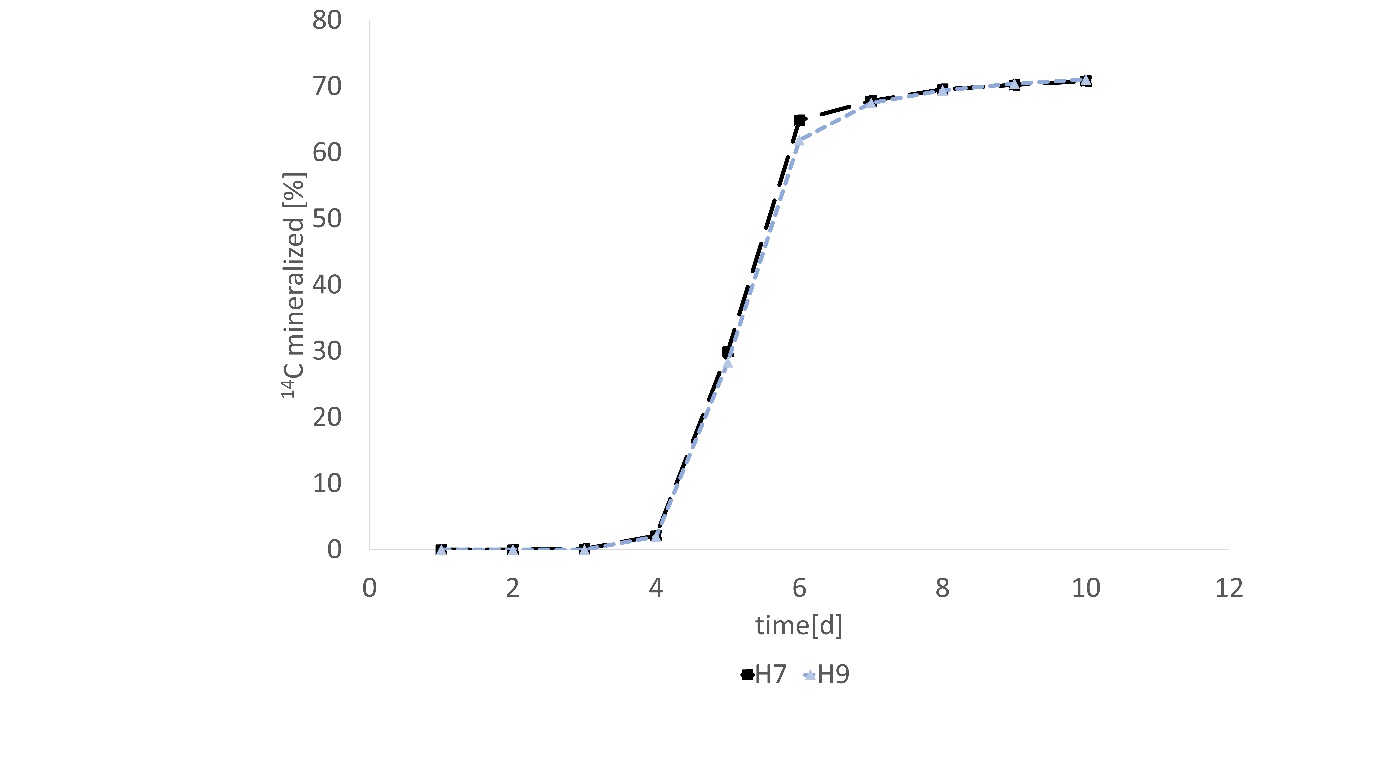

**Figure S3**. Mineralization of ^14^C-linuron by *Variovorax* isolates H7 and H9, isolated from the culture enriched using linuron as sole C-source from the Halen agricultural soil.
